# Locus Coeruleus Stimulation Affects response Adaptation in the Somatosensory Cortex of Whisker-to-barrel Touch System

**DOI:** 10.1101/2021.05.19.444702

**Authors:** Zeinab Fazlali, Yadollah Ranjbar-Slamloo

**Affiliations:** School of Cognitive Sciences, Institute for Research in Fundamental Sciences (IPM), Tehran, Iran

**Keywords:** Sensory Adaptation, Locus Coeruleus, Micro-stimulation, Whisker-to-barrel Touch System, Somatosensory Barrel Cortex

## Abstract

Stimulus-driven responses in the cortex reduce due to prior exposure to sensory stimuli, a phenomenon called sensory adaptation. Depression of synaptic supplies and afterhyperpolarization (AHP) following each action potential are the main proposed mechanisms for adaptation. *In vitro* studies have shown that the neuronal adaptation in the barrel cortex depends on slow AHPs. Such AHPs can be affected by neuromodulators, such as noradrenaline. This evidence suggests that Locus Coeruleus (LC) noradrenergic system may reduce sensory adaptation through this cellular mechanism. We proposed that LC stimulation before whisker deflection can affect the degree of adaptation in the barrel cortex, depending on the nature of noradrenergic interactions in the barrel cortex. We coupled adapted or non-adapted whisker deflections with LC phasic stimulation with a 400 ms interval. A 50ms sinusoidal vibration was applied to the whisker immediately before the test deflection. Neuronal activity was recorded from the barrel cortex (BC) in a urethane anesthetized rat. We quantified the effect of LC stimulation on the degree of adaptation in BC; a lower adaptation index shows lower adaptation. Our result showed that LC stimulation significantly modulated adapted response in 30 % of units with insignificant modulation on the adaptor or non-adapted response. This modulation was in two directions; adaptation decreased in 5 % of units and increased in 25 % of units.

In addition to LC modulation on adaptor response in the level of individual units, adaptor response was lower modulated in around 70 % of units, on average. This modulation was not correlated by LC modulation on non-adapted response. Although sensory adaptation in BC was attenuate by LC stimulation in the majority of units, there was a limited number of units that showed significant modulation.

## Introduction

Adaptation can improve sensory coding efficiently (Adibi et al. 2013) and adjust the dynamic range of neuronal responses to effectively detect changes in the sensory input (Musall et al. 2014). Many cortical cells exhibit spike-frequency adaptation in which action potential frequency decreases with exposure to sustained sensory stimuli. The main mechanisms proposed for adaptation are depression of the synaptic supplies and afterhyperpolarization (AHP) following action potential (Power et al. 2011; Corotto and Michel 2013). Two forms of slow AHP (sAHP) are mediated by potassium currents: sodium-dependent sAHP and calcium-dependent sAHP (Corotto and Michel 2013). The contribution of each type of AHP to neuronal adaptation depends on the brain region being studied. For example in the primary visual cortex, slow contrast adaptation relies on a sodium-dependent sAHP (Sanchez-Vives et al. 2000). In basolateral amygdala, adaptation is generated by a calcium-dependent sAHP (Power et al. 2011). Adaptation in lobster olfactory receptor neurons is mediated by inactivation of Ih current and activation of a calcium-dependent sAHP (Corotto and Michel 2013). Such AHPs can be affected by neuromodulators, such as noradrenaline (Madison and Nicoll 1986; Malenka and Nicoll 1986; Haas and Rose 1987; Foehring et al. 1989; Wallen et al. 1989; Pedarzani and Storm 1996; Khateb et al. 1997).

Earlier studies in the rat barrel cortex showed that adaptation to repetitive stimulation is likely to occur in intracortical or thalamocortical synapses, not at the level of single neurons (Chung et al. 2002; Katz et al. 2006). Due to rapid depression of thalamocortical synapses with repetitive stimulation, neuronal responses in the rat barrel cortex decreased more strongly compared to those in the thalamus (Chung et al. 2002). A later study in slice preparations showed that intrinsic mechanisms in the barrel cortex can contribute to adaptation. Adaptation in layer IV of barrel slice is strongly correlated with the amplitude of calcium-dependent sAHP. The pharmacological blockage of calcium-dependent sAHP decreased spike frequency adaptation *in vitro* (Dıaz-Quesada and Maravall 2008). It is plausible that every neuronal mechanism that targets sAHP may also modulate sensory adaptation to some degree. Therefore, NE can reduce sAHP mediated adaptation by blocking calcium-dependent potassium current (Madison and Nicoll 1986; Foehring et al. 1989).

Here we investigated whether activation of LC-NE system affects sensory adaptation in barrel cortex (BC) neurons. We chose the whisker barrel sensory pathway which is anatomically and functionally well described (Diamond and Arabzadeh 2013; Feldmeyer et al. 2013). We applied phasic LC stimulation (100 and 50 μA) at 400 ms before whisker deflection and recorded from the barrel cortex (BC) in urethane anesthetized rat. To study the interaction of adaptation with LC activation, we applied 50-ms sinusoidal vibration to the whiskers, immediately before the test deflection. We then quantified the interaction of LC stimulation and adaptor on the neuronal response to the test stimulus in BC.

## Material and Methods

### Surgery and electrophysiological recording

Nine adult male Wistar rats, weighing 270-390 g were used. All experiments were approved by the animal care and experimentation committee of the Institute for Research in Fundamental Science (IPM, Iran). Anesthesia was induced by intraperitoneal administration of urethane (1.5 g/Kg), was monitored by the hind paw and corneal reflexes, and maintained stable with supplemental doses of urethane (0.1 g/Kg) if necessary. Two craniotomies were performed on the right hemisphere to provide access to BC (5 × 5 mm; centered at 2.6 mm posterior and 5 mm lateral to Bregma) and LC (4 × 4 mm; centered at 10.8 mm posterior to Bregma and 1.4 mm lateral to the midline). To facilitate access to LC, the rat’s head was tilted down by about 14° (Bouret and Sara 2002).

We recorded from BC while electrically micro-stimulated LC (Figure 1A). BC neuronal activity was acquired with single tungsten microelectrodes (5-10 MΩ, FHC Inc., USA). The principal whisker was determined by the manual stimulation of individual whiskers. We targeted layer 4 neurons based on the depth of penetration from the surface of the exposed dura.

**Figure 1.**
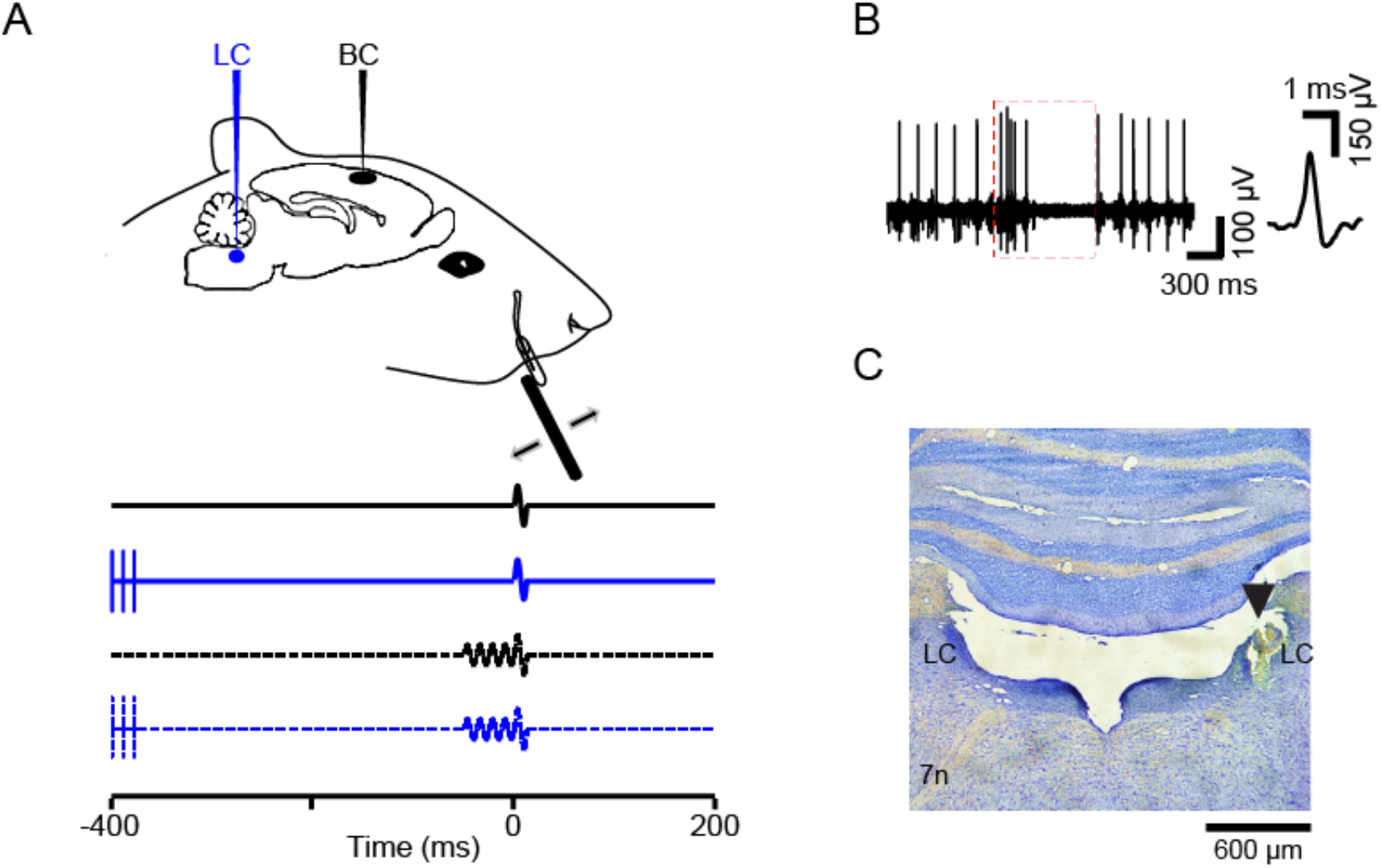
Schematic illustration of the experimental setup. (A) Unit activity was recorded from Barrel Cortex (BC) and Locus Coeruleus (LC) was stimulated. For sensory stimulation a full-cycle sinusoidal deflection was applied to the contralateral whisker. Adaptor is a 50-ms pulse in 80 Hz frequency which was presented before test. LC stimulation was applied 350 ms before adaptor. There are four kinds of trials for each combination. Hereafter, solid and dotted lines respectively represent non-adapted and adapted conditions. LC stimulation pulses are shown as vertical lines. (B) Spiking activity of a representative LC neuron before applying electrical stimulation. Red box represents typical response of a single LC neuron to a paw pinch: the response shows a biphasic characteristic with a brief high frequency spiking followed by a longer suppression. Inset shows one neuron’s spike waveform. (C) Histological verification of the LC recording site for one sample session. The arrowhead shows the site of lesion.

Data acquisition and online amplification were performed using an amplifier (RESANA, Tehran, Iran). During the recording sessions (n=20; 12 with 100 μA and 8 with 50 μA), data were collected at a sampling rate of 30 kHz and filtered online by applying a band-pass filter (300– 6000 Hz) for spiking activity and 0.1 to 6000 for the LFP. Spikes were extracted by offline sorting using principal component analysis implemented in MATLAB (Math Works). In total, 45 units were extracted from BC recordings (25 single- and 20 multi-units) for further analysis. The multi-unit activity in this analysis consisted of single-unit activity.

### Whisker stimulation

Whisker stimulations (a single cycle 80 Hz sinusoidal waveform) were delivered to the principal contra-lateral whisker using a piezoelectric device (RESANA, Tehran, Iran). Stimuli were generated in MATLAB and presented through the analog output of the sound card at a sampling rate of 96 kHz. A lightweight thin piece of glass micropipette was glued to the piezoelectric ceramic. The principal whisker was placed into the micropipette with a 2-3 mm distance from the tip of the micropipette to the base of the whisker. To ensure precise whisker stimulation we used an optic sensor to calibrate the movement range and the waveform of the piezo. We measured the piezo movement to confirm that it accurately followed the voltage command with negligible ringing (average peak amplitude for 12-60 ms after the main deflection was 6% of the main peak, ∼200 Hz). For the range of stimulus intensity (6-60 μm) ringing quickly faded and was not detectable beyond 70-80 ms. To select a near-threshold range of stimuli, at the beginning of each recording session, we applied 10 levels of deflection from a relatively wide range of amplitudes (0-54 μm with 6 μm steps, 50 repetitions). A Nuka-Rushton function was fitted to the average spike count to characterize the neuronal response function. Threshold (T) was defined as the inflection point of this function – i.e. the stimulus amplitude that produced half of the maximum response (M_50_, see Adibi et al., 2013). We applied the test stimuli with 3 amplitudes of 1/2T, T and 2T. The adaptor stimuli consisted of a 50 ms sinusoidal vibration (80 Hz) with three amplitudes of 0, 1/2T, and T immediately before the test stimulus. Each LC-adaptor-stimulus combination was repeated in 5 blocks of 20 trials with pseudorandom order with 300 ms inter-trial interval (end of test deflection to the beginning of the current pulses for the LC). Adaptor-test amplitude combinations consisted of (1) adaptor: 1/2T, test: T, (2) adaptor: T, test: 2T, (3) adaptor: T and test: T and (4) adaptor: 1/2T and test: 2T. These combinations were presented either with or without LC stimulation which was applied 400 ms prior to the test stimulus (Table 1).

**Table 1.** Different test stimulus, adaptor, and LC stimulation trials. Adaptor-test amplitude combinations consisted of: (1) adaptor: 1/2T, test: T, (2) adaptor: T, test: 2T, (3) adaptor: T and test: T and (4) adaptor: 1/2T and test: 2T. These combinations were presented either with or without LC stimulation which was applied 400 ms prior to the test stimulus. Four different red boxes show four combinations. The other white boxes show control conditions for these combinations.

|  |  | LC<br>Stimulation |  | LC<br>Stimulation |  | LC<br>Stimulation |
| --- | --- | --- | --- | --- | --- | --- |
| Stimulus | No Adaptor | No Adaptor | Adaptor<br>$\frac{1}{2}T$ | Adaptor $\frac{1}{2}T$ | Adaptor T | Adaptor T |
| **0** | NoAdap-NoLCStim | NoAdap-LCStim | Adap ($\frac{1}{2}T$)-<br>NoLCStim | Adap ($\frac{1}{2}T$)-<br>LCStim | Adap (T)-<br>NoLCStim | Adap (T)-<br>LCStim |
| **T** | NoAdap-NoLCStim | NoAdap-LCStim | Adap ($\frac{1}{2}T$)-<br>NoLCStim | Adap ($\frac{1}{2}T$)-<br>LCStim | Adap (T)-<br>NoLCStim | Adap (T)-<br>LCStim |
| **2T** | NoAdap-NoLCStim | NoAdap-LCStim | Adap ($\frac{1}{2}T$)-<br>NoLCStim | Adap ($\frac{1}{2}T$)-<br>LCStim | Adap (T)-<br>NoLCStim | Adap (T)-<br>LCStim |

### LC stimulation

LC stimulation parameters were the same as our previous work (Fazlali et al. 2020). Ipsilateral LC was identified for microstimulation by single tungsten microelectrodes (1-2MΩ, FHC Inc., USA) from 5.6-5.9 mm below the dura. To confirm the LC location, we first used the following criteria (Figure 1B): LC neurons normally have broad extracellular spike waveforms (> 0.6 ms), firing rates of 0.1-6 Hz, and respond to paw pinch with a typical excitation-inhibition pattern (red box in Figure 1B) (Cedarbaum and Aghajanian 1978). At the end of the experiment, we verified the LC microstimulation site by histology. To minimize the potential damage to the LC and its projections, no more than two penetrations were made in each recording experiment. LC stimulation consisted of a train of three pulses (0.5 ms pulse duration, 100 Hz, 30-100 µA). A linear stimulus isolator (A395D, World Precision Instruments, Sarasota, FL, USA) was used to apply the current to electrodes. The precise timing and pattern of stimulation was controlled by MATLAB.

### Histology

At the end of each experiment, an electrical lesion was made by passing a DC from a 9-volt battery through the LC electrode tip for 10 s. After transcardial perfusion with ∼300 ml saline (0.9%) followed by ∼300 ml phosphate-buffered formalin (10 %, pH=7.4), the brain was removed and kept in formalin (a minimum of one week) before 10-µm thick coronal sections were made. Sections were Nissl stained and lesions were detected by light microscopy. LC location was compared with the lesion site using the rat brain atlas (Figure 1C, Paxinos, and Watson, 2007). In all cases, the location of recording was anatomically verified to be within LC confirming the reliability of our electrophysiological criteria.

### Data analyses

The sequences of spikes corresponding to trials of the same stimulus were separated and aligned with respect to stimulus onset to generate raster plots (Figure 2A). The occurrence of spiking over time is evaluated by counting the average number of spikes within each bin. A bin size of 10-20 ms is used to generate a prestimulus time histogram (PSTH) for each stimulus. A 50-ms window after the stimulus or adaptor onset was used to quantify neuronal responses as spike counts. The average of the spike counts over trails was then used as a response (R) in adapted and non-adapted conditions of the same stimuli to calculate the adaptation index as follows:

**Figure 2.**
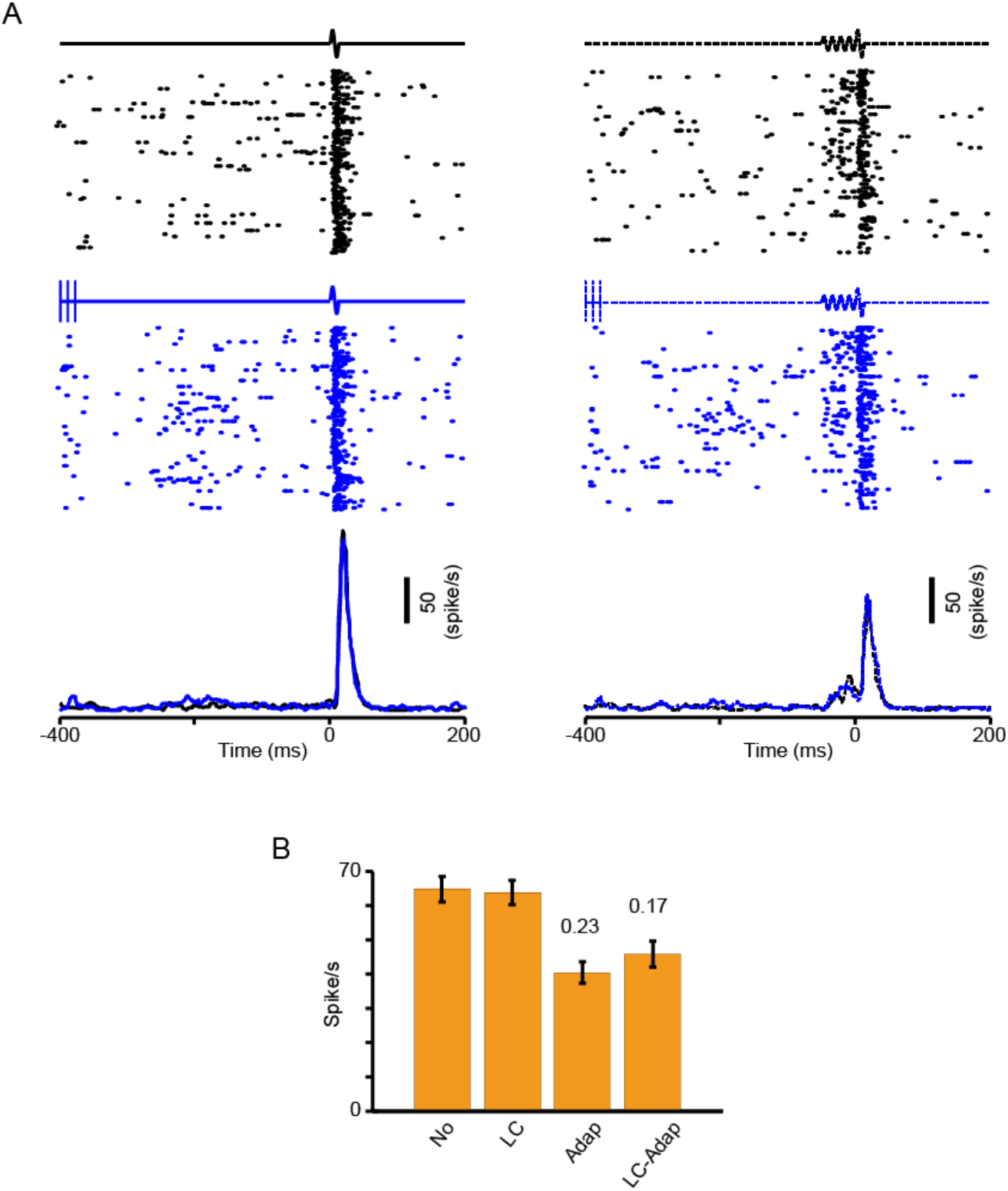
Adaptation attenuated after LC stimulation. (A) Raster plot and PSTH of an example BC unit activity in four conditions; Two non-adapted and two adapted, with or without LC stimulation. Adaptor is T and test is 2T. (B) Average of response in a 50-ms window after test stimulus. Two numbers above two columns show adaptation indices with and without LC stimulation. In this unit adaptation decreased after LC stimulation (P value = 0.10, permutation test).

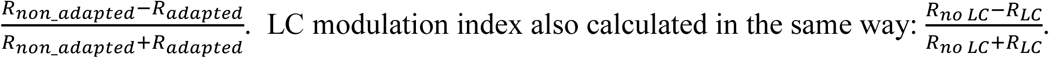

## Results

We coupled whisker adaptor and test stimuli with Locus Coeruleus (LC) stimulation while recording the neuronal activity in the barrel cortex (Figure 1A). LC location was confirmed based on the width of spike waveform, the typical biphasic response profile to noxious stimulation, and histology (Figure 1B and C). BC recording was confirmed based on the neuronal response to brief deflections applied to the neuron’s principal whisker. Each adaptor– test combination was presented 350 ms after LC micro-stimulation when the modulation of the baseline activity was minimal but the evoked activity was maximally modulated (Waterhouse et al. 1998). Adaptor-test amplitude combinations consisted of (1) adaptor: 1/2T, test: T, (2) adaptor: T, test: 2T, (3) adaptor: T and test: T and (4) adaptor: T and test: 2T. These combinations were presented either with or without LC stimulation. From all adaptor-stimulus combinations, we calculated an adaptation index for each neuron (see the Material and Methods). A summary of adaptation indices, LC modulation indices, and adaptation-LC modulation indices for all combinations is given in Supplementary Table 1.

### Effect of LC stimulation and adaptation on BC responses

We first separately quantified the effect of LC stimulation and adaptor on BC responses. LC stimulation significantly increased BC response in 10 % and zero percent of the units in T and 2T test stimulus amplitudes, respectively (2/20 and 0/20, P < 0.05, permutation test). The effect of adaptation on response depends on both intensities of an adaptor and test stimulus. Adaptation significantly decreased response in 35% (7/20), 75% (15/20), 90% (18/20), and 35% (7/20) of units in Combination1, 2, 3, and 4, respectively (P < 0.05, permutation test, Supplementary Table 1). Hence, the adaptor was effective mostly in two combinations when the adaptor was T; Com 2 (adaptor T, test 2T) and Com 3 (adaptor T, test T). Regardless of the combination, if the adaptation index was significant the unit was included for further analyses (see the Material and Methods). In addition to all four conditions, sometimes the result just showed for two conditions with the higher effect of the adaptor; combination 2 and 3.

### Attenuating effect of LC stimulation on adaptation in BC units

Figure 2 shows an example unit in different conditions of adaptation and LC coupling. For this unit the adaptation index in both conditions of LC coupling (LC and no LC stimulation conditions) was significant (0.23 and 0.17, P < 0.05, permutation test). However, the difference between adaptation indices was not significant (P = 0.10, permutation test). In this example, the LC stimulation did not significantly change the neuronal response to either adaptor or the test stimulus in non-adapted condition (P > 0.05, permutation test). Only 5 % of units (1 out of 20) showed a significant attenuating effect of LC stimulation on adaptation, without significantly affecting neuronal activity in non-adapted condition (Supplementary Table 1). There was one unit (5 %) that showed a significant attenuating effect of LC stimulation on adaptation, significantly affecting neuronal activity in non-adapted condition. There was also one unit (5 %) that LC stimulation along with the significant modulation of response to the test stimulus, it also significantly changed the neuronal response to the adaptor.

### Increasing effect of LC stimulation on adaptation in BC units

In contrast, an opposite significant effect of LC stimulation on adaptation index was observed in 25 % of units (5 out of 20) (P < 0.05, permutation test); the adaptation index significantly decreased with insignificant changes in neuronal response to either adaptor or the test stimulus in non-adapted condition (Supplementary Table 1). An example of such an effect is shown in Figure 3.

**Figure 3.**
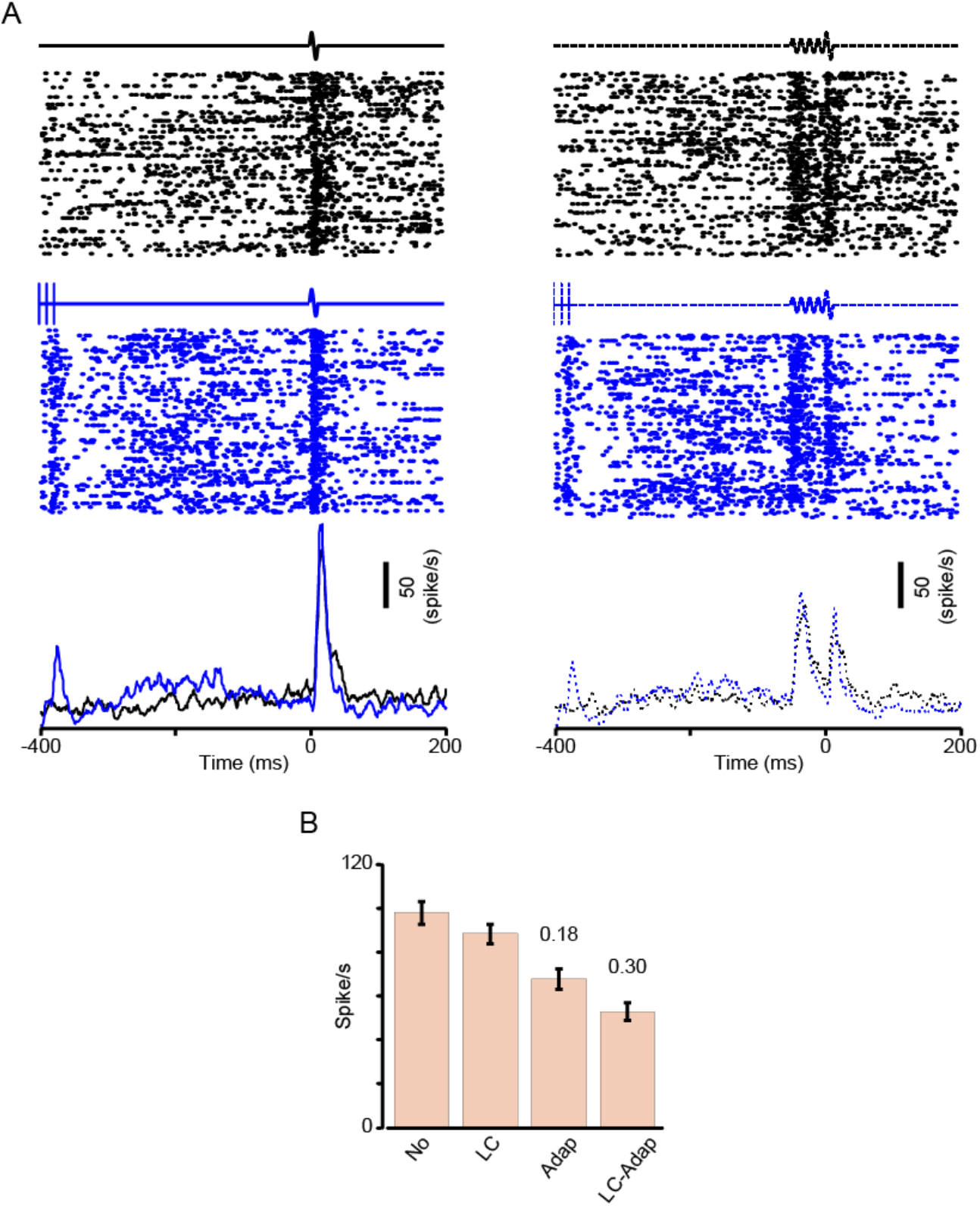
Adaptation increased after LC stimulation. A) Raster plot and PSTH of an example BC unit activity in four conditions. Adaptor is T and test is 2T. (B) Average of response in a 50-ms window after test stimulus. In this unit adaptation significantly increased after LC stimulation (P < 0.05, permutation test). Two numbers above two columns show adaptation indices with and without LC stimulation.

### Effect of LC stimulation, adaptation and both modulation on BC population

After quantifying the modulatory effect of LC and adaptation on each unit, we quantified the effect of both LC and adaptation on BC population. We calculated the average of response in control (no LC and no adaptor), LC, adaptor and LC-adaptor conditions for all four combinations (Figure 4). On average, the adaptor was effective in two combinations, 1 and 2 when the adaptor was at threshold level (T). The highest difference between adapted and non-adapted response was observed in combination 3 when the test stimulus and adaptor was T.

**Figure 4.**
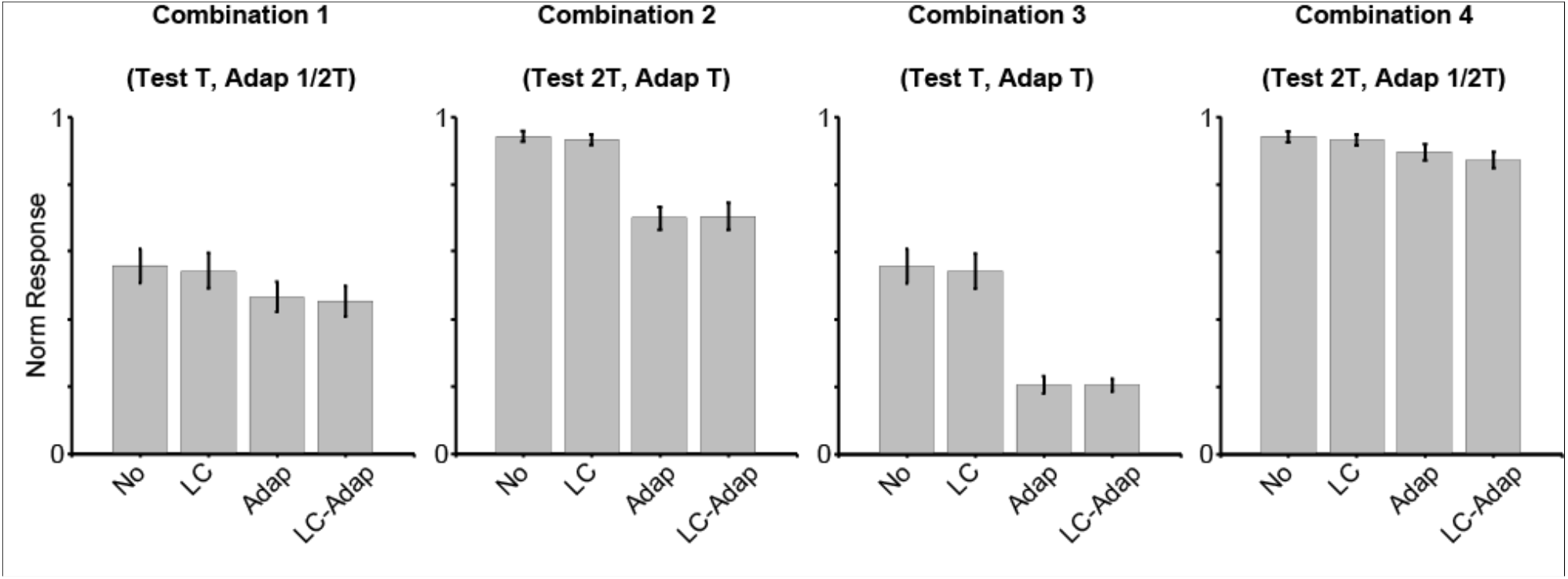
Modulation of response with LC stimulation, adaptation and both LC and adaptation. Average of normalized response in control, LC, adaptor and LC-adaptor conditions for each adaptor-test combination.

Figure 5 shows the interaction of adaptation and LC stimulation for each unit-combination that response significantly modulated by adaptor (n = 50) and separately for each unit-combination in two main combinations (2 and 3) that adaptor significantly decreases an average of response (see also Figure 4). The average of adaptation indices was not changed significantly with LC stimulation neither for all unit-combinations (0.24 ± 0.02 vs 0.23 ± 0.02, n = 50) or for two main combinations (0.30 ± 0.02 vs 0.27 ± 0.03, n = 33) (Figure 5A and B). We then quantified just two combinations with a higher effect of adaptation (Figure 5B). Although the average is not significant, 67 % of data points (22 out of 33) showed a lower adaptation index after LC stimulation compared to the adaptation index without LC stimulation (27 %).

**Figure 5.**
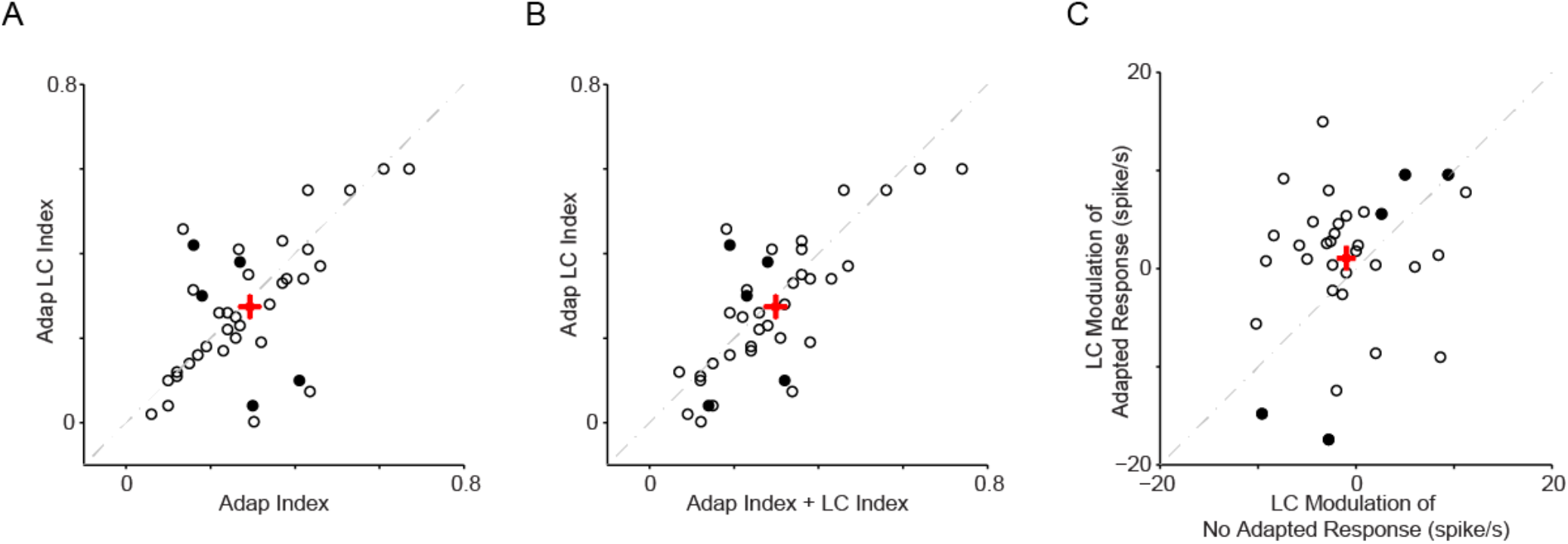
Modulation of LC stimulation on adaptation index. (A) Adaptation Index with and without LC stimulation for units that response significantly modulated by adaptor across four combinations (n = 50). Each data point shows adaptation index for each unit-combination. Red square and its error bars in each direction show average values and standard error of mean (SEM). (B) Same as B but adaptation indices are just for two combinations (2, 3) that average response of units were significantly modulated by adaptor (n = 33), (see Figure 4). (C) Adaptation-LC stimulation Index versus summation of adaptation and LC stimulation index for each unit-combination that significantly modulated by adaptor (n = 50). (D) Same as C but indices are just for two combinations (2, 3) that average response of units were significantly modulated by adaptor (n = 33).

To address the effect of LC stimulation and adaptation separately and their interaction, we summated adaptation indices and LC stimulation indices for each unit and plotted versus adaptation-LC indices. The average of indices of adaptation-LC versus adaptation and LC for all unit-combination were 0.25 ± 0.02 and 0.23 ± 0.02, respectively (n = 50, Figure 5C) and for two main combinations were 0.30 ± 0.03 and 0.27 ± 0.03, respectively (n = 33, Figure 5D). Despite the low difference between the two groups of indices, 70 % of unit-combinations (23 out of 33) showed a lower adaptation-LC index in comparison to the summation of both indices. It further confirmed the interaction of LC stimulation and adaptation.

In addition to calculating the modulation index for LC stimulation, adaptation, and LC-adaptation conditions, we also measured the LC modulation of response in adapted and non-adapted conditions (Figure 6). LC stimulation increased adaptation modulation in 73 % of unit combinations of two effective combinations 2 and 3 (Dots above unity line in Figure 6B). The increasing effect of LC stimulation on adapted response showed that LC affected the occurring of sensory adaptation in BC. We also plotted the LC modulation on adapted and non-adapted responses (Figure 6 C and D). Modulation of adapted response by LC stimulation was higher in 70 % of unit-combinations in comparison to non-adapted response (dots above unity line). We also calculated the correlation between LC modulation of adapted and no-adapted responses and did not found any significant correlation (rho = 0.18, P = 0.31). It showed that in the majority of units the decreasing effect of LC stimulation on sensory adaptation was more than no adaptation condition.

**Figure 6.**
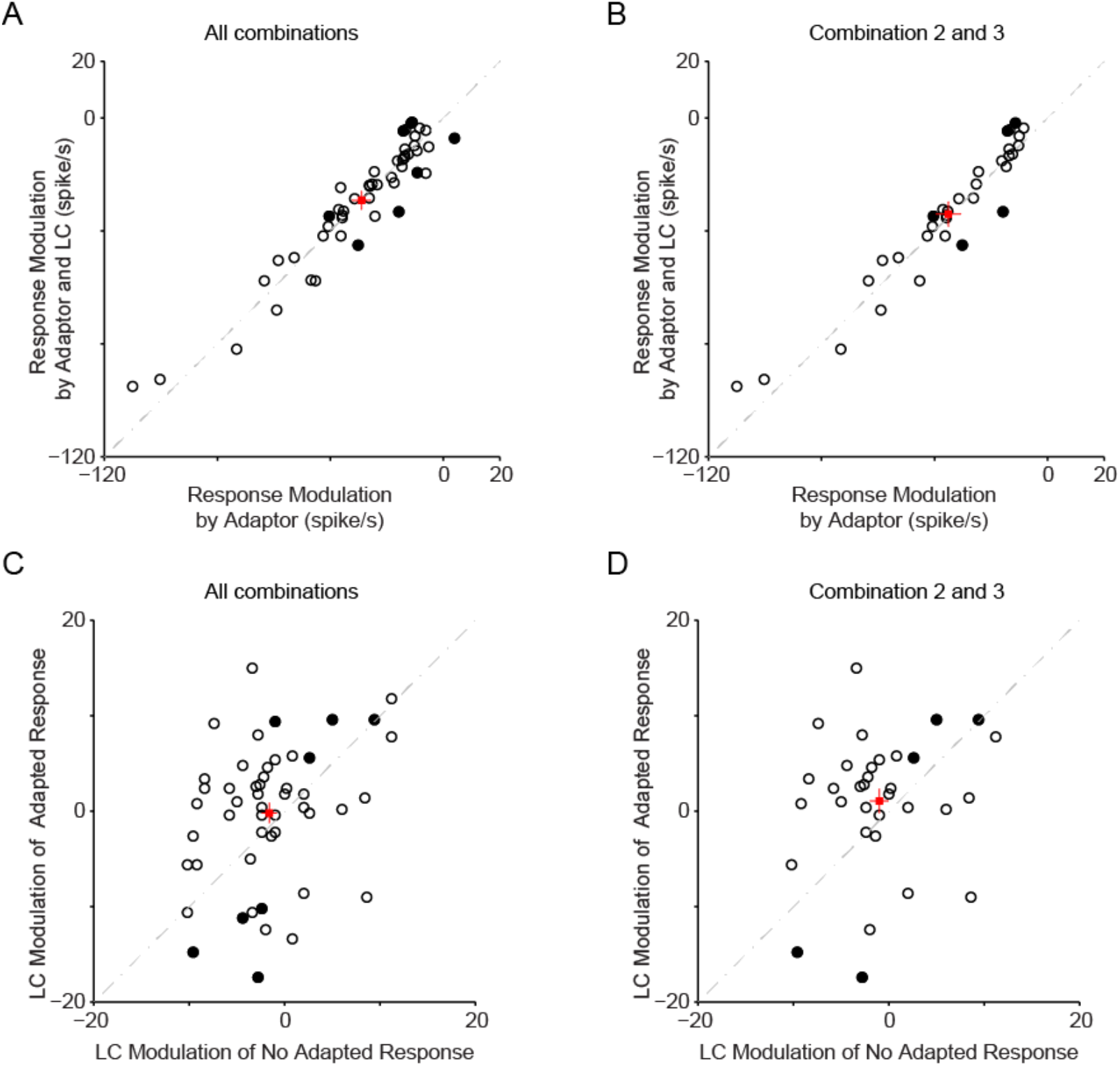
Modulation of BC response by adaptor and LC stimulation. (A) Modulation of response by adaptor versus modulation of response by adaptor and LC stimulation for all combinations (n = 50). Response modulation calculated as an adapted response subtracted by non-adapted response in two axes; with and without LC stimulation. Negative modulations show decrease of response. Solid circles show units with significant difference before and after LC stimulation. These significant units are same as Figure 4. Red square with error bars shows average value. (B) Same as panel A but for two combinations; 2 and 3. (C)Effect of LC stimulation on adapted versus non-adapted response for all combinations. (D) Same as panel C but for combination 2 and 3.

## Conclusion

Our result showed that LC stimulation significantly modulated adapted response in 30 % of units with insignificant modulation on the adaptor or non-adapted response. This modulation was in two directions; adaptation decreased in 5 % of units and increased in 25 % of units. Modulation of adaption by LC stimulation also depends on the combination of test stimulus and adaptor.

Around 70 % of units showed lower adaptation on average (Figure 5 and 6). Therefore, as we expected, adaptation was affected by LC stimulation, and specifically, the majority of units showed lower adaptation after LC stimulation. LC modulation of non-adapted response cannot explain the modulation of adaptation.

Because there was a significant correlation between modulation of response to the adaptor and modulation of adaptation index by LC stimulation (rho=0.35, p=0.014), we need more analyses to quantify the effect of LC stimulation on both the adaptor and test stimulus simultaneously.

## Author Contributions

ZF designed and leaded research; ZF and YR-S performed experiments; ZF analyzed the data; ZF and YR-S interpreted results of experiments; ZF drafted manuscript; ZF and YR-S read and revised the manuscript.

## Data Availability Statement

The data that support the findings of this study are available on request from the corresponding author.

---

**Supplementary Table 1.**
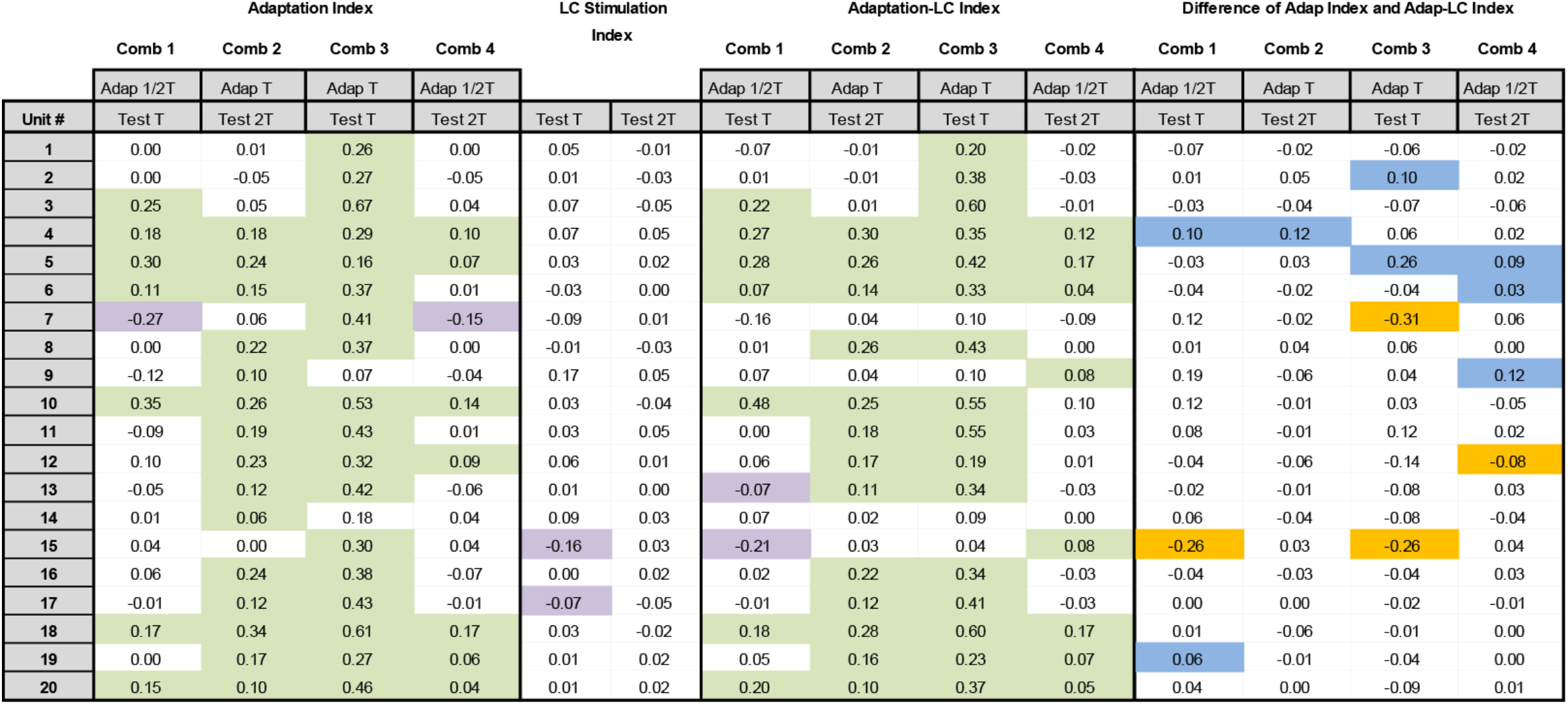
Summary of modulation indices. Green color indices show units that adaptation/LC or both caused significant decrease in response (P < 0.05, permutation test). Purple color indices show units that adaptation/LC or both caused significant increase in response (P < 0.05, permutation test). Orange color boxes show units that difference between Adaptation Index and Adaptation-LC Index were significant and LC decreased Adaptation Index (P < 0.05, permutation test). Blue color boxes show units that difference between Adaptation Index and Adaptation-LC Index were significant and LC increased Adaptation Index (P < 0.05, permutation test).

## References

Adibi M, McDonald JS, Clifford CWG, Arabzadeh E. Adaptation improves neural coding efficiency despite increasing correlations in variability. J Neurosci 33: 2108–20, 2013.

Bouret S, Sara SJ. Locus coeruleus activation modulates firing rate and temporal organization of odour-induced single-cell responses in rat piriform cortex. Eur J Neurosci 16: 2371–2382, 2002.

Cedarbaum JM, Aghajanian GK. Activation of locus coeruleus neurons by peripheral stimuli: modulation by a collateral inhibitory mechanism. Life Sci 23: 1383–1392, 1978.

Chung S, Li X, Nelson SB. Short-term depression at thalamocortical synapses contributes to rapid adaptation of cortical sensory responses in vivo. Neuron 34: 437–46, 2002.

Corotto FS, Michel WC. Mechanisms of Afterhyperpolarization in Lobster Olfactory Receptor Neurons Mechanisms of Afterhyperpolarization in Lobster Olfactory Receptor Neurons..

Diamond ME, Arabzadeh E. Whisker sensory system – From receptor to decision. Prog Neurobiol 103: 28–40, 2013.

Dıaz-Quesada M, Maravall M. Intrinsic Mechanisms for Adaptive Gain Rescaling in. J Neurosci 28: 696–710, 2008.

Fazlali Z, Ranjbar-Slamloo Y, Arabzadeh E. Modulation of Sensory Response at Different Time Lags after Locus Coeruleus Micro-stimulation. bioRxiv (January 1, 2020). doi: 10.1101/2020.07.09.188615.

Feldmeyer D, Brecht M, Helmchen F, Petersen CCH, Poulet JFA, Staiger JF, Luhmann HJ, Schwarz C. Barrel cortex function. Prog Neurobiol 103: 3–27, 2013.

Foehring RC, Schwindt PC, Crill WE. Norepinephrine selectively reduces slow Ca2+- and Na+-mediated K+ currents in cat neocortical neurons. J. Neurophysiol. 61: 245–56, 1989.

Haas HL, Rose GM. Noradrenaline blocks potassium conductance in rat dentate granule cells in vitro. Neurosci Lett 78: 171–174, 1987.

Katz Y, Heiss JE, Lampl I. Cross-whisker adaptation of neurons in the rat barrel cortex. J Neurosci 26: 13363–72, 2006.

Khateb A, Fort P, Williams S, Serafin M, Jones BE, M??hlethaler M. Modulation of cholinergic nucleus basalis neurons by acetylcholine and n-methyl-d-aspartate. Neuroscience 81: 47–55, 1997.

Madison BYD V, Nicoll RA. Physid. (1986), 372,..

Malenka RC, Nicoll RA. Dopamine decreases the calcium-activated afterhyperpolarization in hippocampal CA1 pyramidal cells. Brain Res 379: 210–215, 1986.

Musall S, Behrens W Von Der, Mayrhofer JM, Weber B, Helmchen F, Haiss F. Tactile frequency discrimination is enhanced by circumventing neocortical adaptation. Nat Neurosci 17: 1567–1573, 2014.

Paxinos G, Watson C. The Rat Brain in Stereotaxic Coordinates. Academic, San Diego, USA, 2007.

Pedarzani P, Storm JF. Evidence that Ca/calmodulin-dependent protein kinase mediates the modulation of the Ca2+-dependent K+ current, IAHP, by acetylcholine, but not by glutamate, in hippocampal neurons. Pflugers Arch 431: 723–8, 1996.

Power JM, Bocklisch C, Curby P, Sah P. Location and function of the slow afterhyperpolarization channels in the basolateral amygdala. J Neurosci 31: 526–37, 2011.

Sanchez-Vives M V, Nowak LG, McCormick D a. Membrane mechanisms underlying contrast adaptation in cat area 17 in vivo. J Neurosci 20: 4267–85, 2000.

Wallen P, Buchanan JT, Grillner S, Hill RH, Christenson J, Hokfelt T. Effects of 5-hydroxytryptamine on the afterhyperpolarization, spike frequency regulation, and oscillatory membrane properties in lamprey spinal cord neurons. J Neurophysiol 61: 759–768, 1989.

Waterhouse BD, Moises HC, Woodward DJ. Phasic activation of the locus coeruleus enhances responses of primary sensory cortical neurons to peripheral receptive field stimulation. Brain Res 790: 33–44, 1998.

